# Phospholipase Cβ2 Promotes Vascular Endothelial Growth Factor Induced Vascular Permeability

**DOI:** 10.1101/2022.03.09.483667

**Authors:** Kathryn N. Phoenix, Zhichao Yue, Lixia Yue, Chunxia G. Cronin, Bruce T. Liang, Luke H. Hoeppner, Kevin P. Claffey

## Abstract

**Background:** Regulation of vascular permeability (VP) is critical to maintaining tissue metabolic homeostasis. Vascular endothelial growth factor (VEGF) is a key stimulus of VP in acute and chronic diseases including ischemia reperfusion injury, sepsis and cancer. Identification of novel regulators of VP would allow for the development of effective targeted therapeutics for patients with unmet medical need.

**Methods:** In vitro and in vivo models of VEGFA-induced vascular permeability, pathological permeability, quantitation of intracellular calcium release and cell entry, and PIP2 levels were evaluated with and without modulation of PLCβ2.

**Results:** Global knock-out of PLCβ2 in mice resulted in blockade of VEGFA-induced vascular permeability in vivo and trans-endothelial permeability in primary lung endothelial cells. Further work in an immortalized human microvascular cell line modulated with stable knock-down of PLCβ2 recapitulated the observations in the mouse model and primary cell assays. Additionally, loss of PLCβ2 limited both intracellular release and extracellular entry of calcium following VEGF stimulation as well as reduced basal and VEGFA-stimulated levels of PIP2 compared to control cells. Finally, loss of PLCβ2 in both a hyperoxia induced lung permeability model and a cardiac ischemia:reperfusion model resulted in improved animal outcomes when compared to WT controls.

**Conclusions:** The results implicate PLCβ2 as a key positive regulator of VEGF-induced VP through regulation of both calcium flux and PIP2 levels at the cellular level. Targeting of PLCβ2 in a therapeutic setting may provide a novel approach to regulating vascular permeability in patients.

**Graphic Abstract:** 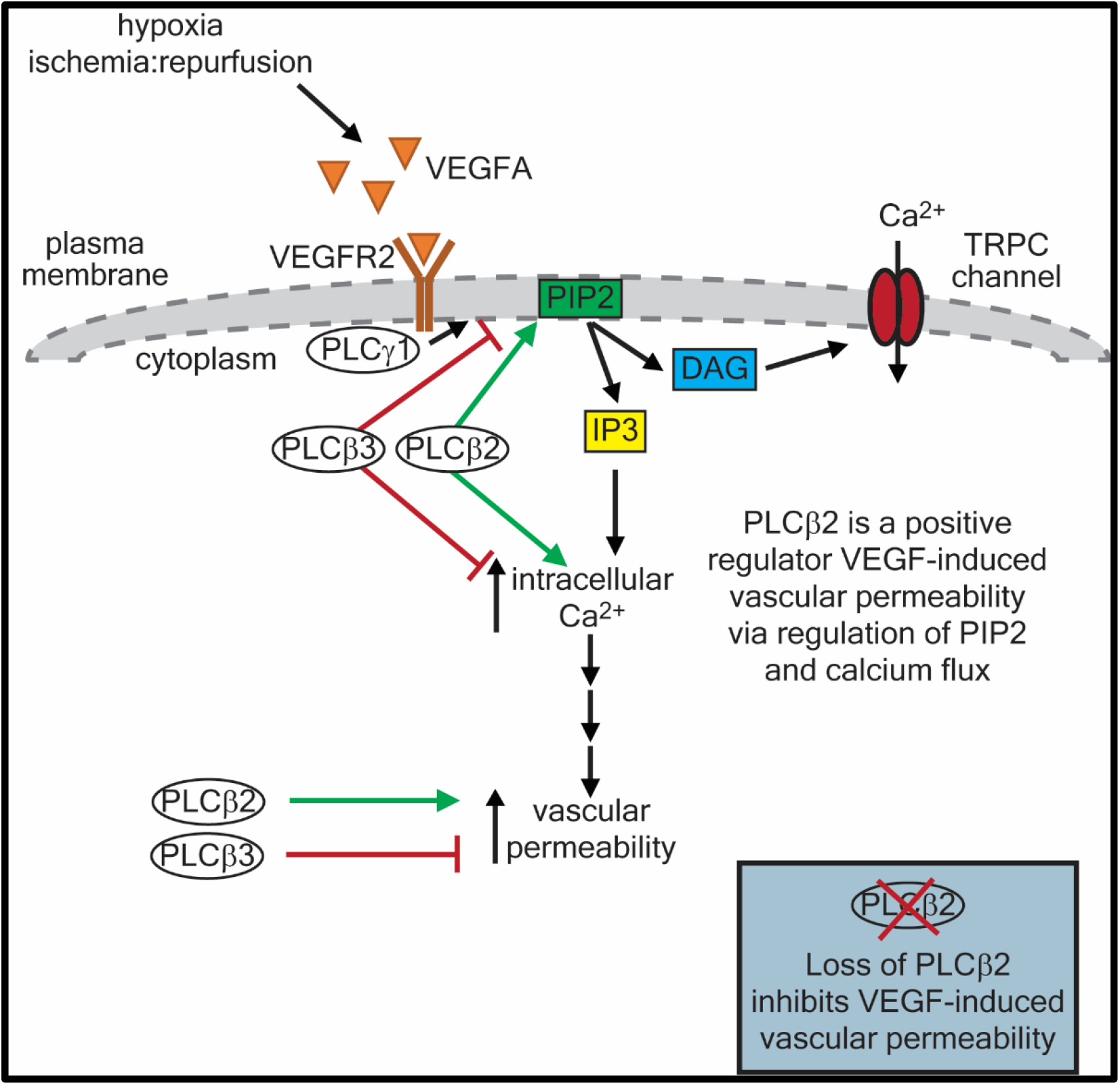

**Highlights:**

- PLCβ2 promotes VEGFA induced vascular permeability.
- Loss of PLCβ2 prevents VEGFA vascular permeability via repression of cellular calcium flux and membrane PIP2 levels.
- Loss of PLCβ2 reduces vascular permeability and improves outcomes in a hyperoxic lung damage model and a cardiac ischemia:reperfusion model in vivo.
- Targeting PLCβ2 inhibition may lead to a novel therapeutic for diseases such as stroke and myocardial infarction.

## Introduction

Vascular permeability (VP) is essential for the maintenance of normal tissue and organ function. Tight regulation of VP allows nutrient transport and immune surveillance without excessive leakage of plasma into the tissues under normal physiological conditions.^1,2^ This regulation maintains a low level of VP during homeostasis but can be significantly impacted by pathological disease conditions including wound healing, cancer, myocardial infarction, and stroke. Dysregulation of VP results in hyper-vascular permeability which can significantly contribute to morbidity and mortality associated with these conditions.^3^ In myocardial infarction and stroke, vascular permeability induced by factors secreted from ischemic tissue expands the area of tissue damage by several fold.^2,4^ In the case of lung injury caused by infection or ventilator-induced over inflation, accumulation of fluid within the alveolae reduces oxygen transport and is a major contributor to mortality.^5,6^ Identification of novel regulators of VP may allow for the development of effective targeted therapeutics for a number of diseases and patients with unmet medical need.

Vascular endothelial growth factor (VEGF) was initially discovered in the 1980s as a tumor secreted factor that strongly promoted microvascular permeability and was named vascular permeability factor (VPF) before its subsequent identification as VEGF, an endothelial mitogen essential for the development of blood vessels.^7–9^ VEGF, specifically VEGFA, belongs to a family of proteins including VEGFB, VEGFC, VEGFD, VEGFE and placental growth factor. VEGFC and VEGFD are key regulators of lymphangiogenesis, while VEGFA is known to be the is dominant regulator of angiogenesis and vascular permeability.^3,8,10–13^ VEGFA is involved in the stimulation of acute vascular permeability under conditions of tissue hypoxia caused by inadequate vascular flow as observed in conditions such as myocardial infarction and cerebral stroke. In addition, where tissue demand outpaces vascular supply such as in rapidly growing solid tumors or high skeletal muscle demand, VEGFA is potently expressed, and pathological permeability and vascular barrier dysfunction can occur. ^12,14,15^

VEGFA has been shown to promote endothelial cell proliferation, migration and vessel formation in both in in vitro and in vivo model systems.^1,3,12^ VEGFA expression is stimulated by low oxygen conditions, during hypoxia and ischemia, in a hypoxia-inducible factor (HIF) dependent manner. Alternatively, VEGFA can be stimulated by other factors including FGF-2, TNF-alpha, and shear stress.^12,16–19^ VEGFA induces microvascular permeability by multiple mechanisms including junctional remodeling between cells, the formation of fenestrae and induction of vesiculo-vacuolar organelles (VVO).^20^ VEGFA primarily mediates its function through 2 receptor tyrosine kinase receptors, VEGFR1 and VEGFR2, along with neuropilin as a coreceptor. VEGFA promotes vascular permeability by binding to VEGF receptor 2 (VEGFR2) to stimulate receptor clustering and activation of downstream signaling.^12,21^ Several pathways have been implicated in regulating VP mechanisms. These include phospholipase C (PLC) dependent intracellular calcium release, src kinase-mediated phosphorylation and internalization of junctional proteins, Rho GTPase activation, cytoskeletal rearrangement, and eNOS signaling.^12,13,21–23^

PLC is a crucial enzyme in phosphoinositide metabolism as it hydrolyses phosphatidylinositol 4,5-bis-phosphate (PIP2) to generate two second messengers, inositol 1,4,5-trisphosphate (IP3) and diacylglycerol (DAG), that drive a variety of different cellular responses.^24^ There are 13 mammalian PLC isozymes, including PLCβ (β1, β2, β3, and β4) and PLCγ (γ1 and γ2), that are regulated by distinct mechanisms. Known activators of PLC include Ca^2+^, heterotrimeric G proteins, small G proteins, and receptor/non-receptor tyrosine kinases.^25,26^ PLC isozymes are responsible for a wide array of physiological functions. Regarding the role of PLC isozymes in the regulation of vascular function, PLCγ1 regulates VEGFA-mediated endothelial cell migration, positively controls vascular permeability induced by VEGFA, and plays a key role in arterial development.^27^ It has been demonstrated that in endothelial cells both PLCγ and PLCβ3 can increase PIP2 hydrolysis significantly in response to VEGFA stimulation.^1,28,29^ PLCβ3 opposes the function of PLCγ1 by negatively regulating VEGFA-mediated vascular permeability.^30^ However, PLCβ3 and PLCγ1 rely on a similar intracellular Ca^2+^-dependent mechanism to differentially control endothelial barrier integrity. Little is known about whether other PLC isozymes beyond PLCβ3 and PLCγ1 regulate vascular permeability.

This study presents a novel regulator of VP in the PLC family that may be an interesting new target to consider for anti-VP therapeutic strategies. In this work, the role of an additional PLC beta-isoform, PLCβ2, was investigated in VEGFA-induced permeability. Results from experiments using PLCβ2 knock-down cell lines and PLCβ2 null-mice demonstrate that PLCβ2 acts as a positive regulator of VP. Specifically, PLCβ2 acts as a positive regulator of specific endothelial cell and vascular responses to VEGFA and loss of PLCβ2 leads to prevention of VEGFA-induced vascular permeability through a reduction of PIP2 levels and calcium flux in vascular endothelial cells.

## Materials and Methods

### Cell Lines and Reagents

TIME cells (CRL-4025) were purchased from American Type Culture Collection (Manassas, VA). Cells were maintained at 37°C in 5% CO_2_ in either EGM-2 MV complete media (CC-3125, Lonza, Walkersville, MD) or in M199 supplemented with 20% FBS (Atlanta Biologicals, Flowery Branch, GA), 1% Pen/Strep (Life Technologies, Carlsbad, CA) and 1% endothelial cell growth supplement (ECGS) (Yale University, Vascular Biology & Therapeutics Program Core Facility, New Haven, CT).

### Animals

C57Bl/6 mice with genetic alteration in the PLC-β2 and PLC-β3 isoforms were previously described^31,32^ and obtained from Dr. Dianqing Wu, Yale University, New Haven, CT.^20–22^ Wild Type C57Bl/6 mice were obtained from The Jackson Laboratory (Stock No: 000664). Male and female mice, 8-12 weeks of age were used for all studies. All animal experiments were approved by the University of Connecticut Health Center’s Institutional Animal Care and Use Committee.

### VEGFA-induced Microvascular Permeability

In vivo peripheral permeability assays were completed as described previously.^30,33^ Briefly, wild-type, PLCβ2-null and PLCβ3-null mice were anesthetized with 2% Avertin (0.5 mL/20 g) via intraperitoneal injection and 100 μL of FITC-dextran (5 mg/mL, 2000 kDa, 52471, Sigma, St. Louis, MO) injected intravenously via retro-orbital injection. Animals were placed on a Kodak Multi-modal Imager (2000MM) with warm water bottles to maintain body temperature and the central vessels in the ears were imaged. Saline as vehicle control or mouse VEGFA, 10 ng/mL, (H7125/CF, Sigma, St. Louis, MO) was injected intradermally into the middle of the ear using a 30-gauge needle (30 μL). Ear vasculature was then imaged with fixed exposure times (30 seconds) in a stationary position for 45 minutes after injection (465nm ex/535nm em). Fluorescence images were then quantified using Kodak MI software.

### Basal Permeability

Basal permeability of WT and PLC knock-out mice was assessed as previously described.^33^ Briefly, 100μl Evans Blue Dye (EBD) (E2129, Sigma, St. Louis, MO) was injected intravenously via retro-orbital injection (0.5% EBD in PBS) into mice and allowed to circulate for four hours. Vascular permeability was determined by quantitative measurement of the dye incorporated per milligram of tissue (dry weight) as describe previously.^34^ Briefly, EBD was extracted from dried tissues by incubating tissues in formamide at 65°C overnight. Optical density of extracted dye was measured at 620nm absorbance on a Synergy Mx spectrophotometer (BioTek Instruments, Inc., Winooski, VT), and concentration calculated by comparison to EBD standard curve and values normalized to tissue dry weight.

### Primary Murine Lung Endothelial Cell Isolation

Primary murine lung endothelial cells (MLECs) were isolated using the Lung Dissociation Kit (130-095-927) and Endothelial Cell isolation protocol using CD31 MicroBeads (130-097-418) according to manufacturer’s protocol (Miltenyi Biotec, Inc. Auburn, CA). After isolation cells were plated onto fibronectin coated dishes and used in assays between day 4 and day 6 post-isolation as detailed below. MLECs were cultured in EGM-2 MV media (Lonza, Walkersville, MD).

### Transwell Endothelial Permeability Assay

Primary MLECs were plated and grown for up to 2 days until they formed a confluent monolayer on 0.4 uM Transwell inserts (3413, Costar, Cambridge, MA, USA). Medium containing 0.5 mg/ml FITC-dextran (2000 kDa) was loaded in the upper compartment of the Transwell and cells stimulated with control (PBS), VEGFA (R&D Systems) or Histamine (H7125, Sigma, St. Louis, MO). The amount of FITC-dextran diffused through the endothelial monolayer into the lower compartment was measured by a microplate reader (495nM ex/519nm em) (Syngery Mx, BioTek Instruments, Inc., Winooski, VT) and values determined via comparison with a standard curve.

### shRNA Transduction and Cell Selection

TIME cells were transduced using Mission shRNA lentiviral particles targeting GFP (SHC002V), PLCβ2 (TRCN0000002136) and PLCβ3 (TRCN0000000433) (Sigma, St. Louis, MO) with 8 ug/ml hexadimethrine bromide (H9268, Sigma) overnight. Cells were washed with PBS and allowed to recover for 24 hours prior to selection with 2 ug/ml puromycin. Stable shRNA knock-down cell lines were established. Transduction and selection were repeated at least three times for each gene target.

### RNA Isolation and quantitative RT-PCR

Total cell RNA was isolated using the RNeasy Kit (Qiagen) according to the manufacturer’s protocol. One microgram of RNA was used to produce cDNA using the iScript cDNA Synthesis Kit (1708890, BioRad). Quantitative PCR primers were designed using ABI Primer Express software for use with the iQ Syber Green Supermix (1708880, BioRad) and the MyIQ qPCR system (BioRad). Primers used for qPCR: PLCβ2 - Forward Sequence: GGCCGAGCAAATCTCCAAAA; Reverse Sequence: TCTTTGGTGTCGTTCTCCGA; PLCβ3 - Forward Sequence: TCAACCGGAGTGAGTCCATC; Reverse Sequence: GGATCTTGTCAATGTCCGGC; RPLP2 - Forward Sequence: TCCTCGTGGAAGTGACATCGT; Reverse Sequence: CTGTCTTCCCTGGGCATC

### Multi-well Calcium Assays

Multi-well calcium flux assays were performed with the Fura-2 QBT Calcium Kit (R8197, Molecular Devices, Inc., Sunnyvale, CA). 10,000 – 20,000 cells/well were plated into a 96-well plate to achieve confluence when analyzed. Prior to the assay, cells were loaded for 1 hour with Fura-2 QBT loading dye according to the manufacturer’s protocol. Baseline signals were obtained on a Synergy Mx spectrophotometer (BioTek Instruments, Inc., Winooski, VT) at 340 ex and 510 em (calcium bound) and 380 ex and 510 em (calcium unbound) for one minute prior to exogenous stimulation with with control (PBS), VEGFA (R&D Systems) or Histamine (H7125, Sigma, St. Louis, MO). Kinetic reads were completed every 23 seconds for up to 10 minutes post-stimulation. Data was normalized to baseline values for each well.

### Calcium Release/Entry Imaging Assay

Calcium release imaging assays were performed as previously reported.^35^ Briefly, cells plated on glass bottom cell culture dish were loaded with 10 μmol/L Fura-2 acetoxymethyl ester (Molecular Probes) and 0.1% pluronic F-127 (Sigma, St. Louis, MO) in Tyrode solution for 45 minutes in a humidified 5% CO2 incubator at 37°C. Non-incorporated dye was washed away with Tyrode solution containing NaCl 148mmol/L, KCl 5mmol/L, CaCl_2_ 2mmol/L, MgCl_2_ 1mmol/L glucose 10 mmol/L, HEPES 10 mmol/L, pH 7.4. After 30 seconds perfusion with calcium free solution (NaCl 148mmol/L, KCl 5mmol/L, MgCl_2_ 1mmol/L glucose 10 mmol/L, HEPES 10 mmol/L, pH 7.4), Ca^2+^ transients were evoked by the treatment with agonist (VEGFA) in calcium free solution or Tyrode solution for up to 10 minutes. Ionomycin (Iono) at 1μmol/L was used as an internal positive control. Fluorescence intensities at 510 nm with 340 nm and 380 nm excitation were collected at a rate of 1 Hz using Cool SNAP HQ2 (Photometrics) and data were analyzed using NIS-Elements (Nikon). Cytosolic Ca^2+^ was measured by the comparing the ratio of fluorescence intensity at 340 nm and 380 nm (F340/F380) from at least 10 dividual cells in primary cell assays and 24 individual cells in stable cell assays per condition. Normalized average cell response is reported.

### PIP2 ELISA

PIP2 extraction and quantification via ELISA was performed according to the manufacturer’s directions (K-4500, Echelon Biosciences, Inc., Salt Lake City, UT).

### Hyperoxia Induced Lung Injury Model

Animals were exposed for hyperoxic conditions for 72 hours (70% medical grade oxygen, 30% medical grade air, ~76% O_2_/24% N_2_). After treatment, animals were moved to room air for 4 hours prior to harvest. Thirty minutes prior to sacrifice animals were anesthetized via intraperitoneal injection with 2% Avertin (0.5 mL/20 g) and 100 μL of FITC-dextran (5 mg/mL, 2000 kDa) was injected intravenously via retro-orbital injection. After mice were euthanized, lungs were lavaged with 1 ml of sterile PBS to evaluate lung vasculature permeability via microplate reader (495nM ex/519nm em) (Syngery Mx, BioTek Instruments, Inc., Winooski, VT) and tissues taken for histological analysis. Tissues were fixed in 10% formalin, processed and paraffin embedded via standard histology protocols. FFPE sections (4 μm) were cut and H&E stained for imaging and quantification with ImageProPlus software (Media Cybernetics, Inc. Rockville, MD).

### Cardiac Ischemia:Reperfusion Model

Cardiac ischemia was induced by temporary LAD ligation according to the established procedure.^36^ Anesthesia was induced by intraperitoneal injection of ketamine (100 mg/kg) and xylazine (10 mg/kg). Following 30 mins of ischemia, sutures were removed to reperfuse tissues. A portion of animals in this study (total n=4-6 per strain per independent study) were injected intravenously with FITC-dextran (2000 kDa) during the 30-minute reperfusion. The remaining animals (total n=4-6 per strain per independent study) were allowed to recover for 10 weeks prior to euthanasia and tissue harvest and processing. Infarct size was quantified as previously described.^36–38^ Following tissue fixation in 10% formalin, the left ventricle was cut into five 1.0 mm-thick transverse sections with a sterile scalpel from the apex toward the base. Sections were embedded in paraffin, cut into 4 μm sections, and stained with Masson’s trichrome to measure area of fibrosis (infarcted myocardium). The lengths of the infarcted and non-infarcted endocardial and epicardial surfaces were traced with a planimeter image analyzer (ImageProPlus). Infarct size was calculated as the ratio of infarct length to the circumference of both the endocardium and the epicardium.

### Statistical Analysis

Data from individual experiments were represented as mean ± standard error unless otherwise stated. Sample number (n) per experimental group is noted in the figure legend. Experiments were repeated at least three times unless otherwise stated. Statistical comparison of groups was performed using 2-tailed student t-test or ANOVA test with appropriate tests for equal variances. Statistical significance was defined and indicated as *P* ≤ 0.05 (*) or *P* ≤ 0.01 (**).

## Results

### PLCβ2-null animals are resistant to VEGFA-induced microvascular permeability

VEGFA-induced microvascular permeability can be significantly modulated by the PLCβ3 isoform. Global deletion of the PLCβ3 from mice results in a significant increase of dermal vascular permeability in vivo after exposure to VEGFA, suggesting that PLCβ3 negatively regulates VEGFA-induced vascular permeability.^30^ In order to determine whether there was any role for the PLCβ2 isoform in this model of VEGFA-induced microvascular permeability, PLCβ2-null mice were evaluated in an in vivo peripheral model of VEGFA-induced microvascular permeability, Figure 1. Wild type (WT), PLCβ2-null mice and PLCβ3-null mice were injected intravenously with a large molecular weight FITC-dextran and then intradermal injections were performed with VEGFA or PBS control into the ears. Consistent with previously published results^30,33^, WT mice displayed a significant, 1.5-fold, increase in microvascular permeability when injected with VEGFA compared to PBS control injections, Figure 1A. Additionally, PLCβ3 null animals displayed an average 4-fold increase in microvascular permeability when treated with VEGFA that was significantly higher than the WT mice, Figure 1C, as previously shown.^30^ Finally, when PLCβ2 null mice were injected with VEGFA there was no significant change in permeability compared to the PBS control, Figure 1B. Maximum induced signal from each condition was averaged and compared between groups. WT and PLCβ3-null mice displayed significant increase in the maximum signal when treated with VEGFA (1.12±0.07 vs 1.77±0.18, *P*=0.011 and 1.67±0.18 vs. 3.78±0.70, *P*=0.012, respectively), while PLCβ2-null animals showed no detectable change in signal intensity compared to PBS control (1.08±0.02 vs. 1.14±0.03, *P*=0.15). Interestingly, in PLCβ3-null mice the permeability in PBS control samples was significantly higher than both the WT and the PLCβ2-null PBS signals (1.12±0.07 vs. 1.67±0.18, *P*=0.043) suggesting a possibility that there could be a difference in basal vascular permeability in these animals.

**Figure 1.**
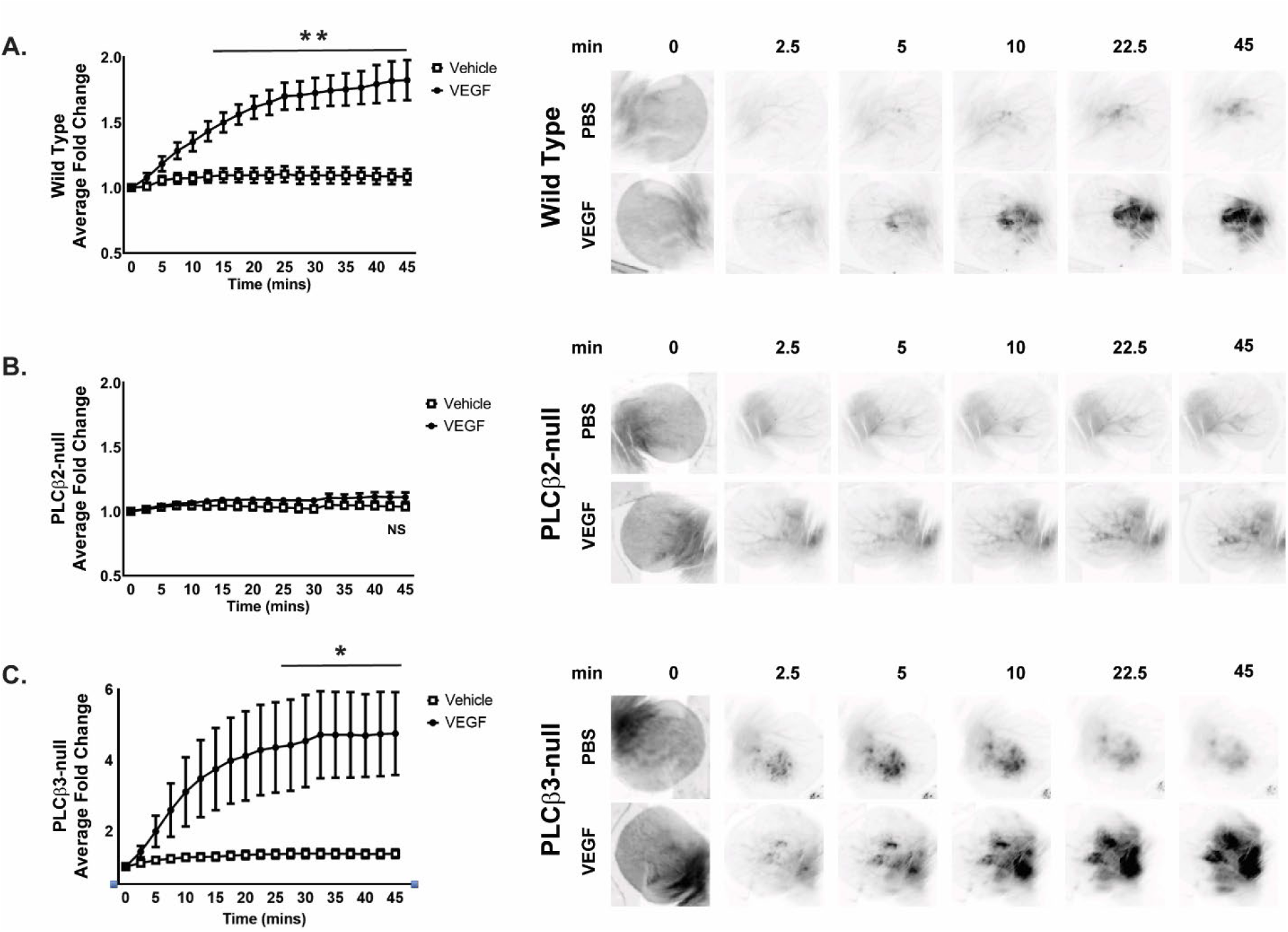
VEGFA-induced dermal vascular permeability is reduced by loss of PLCβ2. **Wild-Type, PLCβ2-null and PLCβ3-null** animals were injected IV with FITC-dextran then exposed locally to VEGFA (closed circles) or PBS control (open squares) via intradermal injection into the ears. Animals were imaged over 45 mins for FITC-dextran extravasation and quantified. **A.** Wild Type. (n = 7) **B.** PLCβ2-null. (n = 7) **C.** PLCβ3-null. (n=8). (* = *P* ≤ 0.05; ** = *P* ≤ 0.01)

To evaluate the basal vascular permeability levels of these animals, Evans blue dye was injected intravenously into WT, PLCβ2-null and PLCβ3-null animals and allowed to circulate for four hours. At harvest, various organs were removed, weighed, and dried overnight. Dried tissues were weighed, dye extracted with formamide and evaluated at 740 nm. In the eight different tissues that were evaluated (lung, bladder, heart, kidney, ear, spleen, adipose and liver), there were differences in basal permeability between tissue type. However, no significant differences were observed between WT and PLCβ2-null or PLCβ3-null animals in any tissue, Supplemental Figure 1A. These data suggest that while there are significant differences in VEGFA-induced peripheral microvascular permeability in animals with different PLC beta isoforms, there were no detectable differences in basal organ permeability in the same mice.

### PLCβ2-null primary endothelial cells are resistant to VEGFA-induced permeability

To further evaluate endothelial cell permeability response to VEGFA, primary endothelial cells were isolated from the lungs of WT, PLCβ2-null and PLCβ3-null mice. Primary murine lung endothelial cells (MLECs) were evaluated for their response to VEGFA in a transwell permeability assay in vitro. Monolayers of MLECs were grown to confluence on transwell inserts and the top compartment loaded with large molecular weight FITC-dextran (2000 kDa). WT and PLCβ3-null cells both demonstrated significant increases in the amount of FITC-dextran that was detected in the bottom chamber after treatment with VEGFA, Figure 2A. In contrast, the PLCβ2-null MLECs did not respond significantly to VEGFA treatment. An alternative stimulus that promotes vascular permeability was also evaluated in the transwell assay. Histamine functions through the histamine H1 receptor. The histamine H1 receptor is a G_αq/11_ -dependent GPCR in endothelial cells that results in PLCβ activation, calcium release and endothelial permeability.^39–41^ Interestingly, all three MLEC cell types responded equally to a different permeability agonist, histamine. These data are consistent with the in vivo observations and further suggests that PLCβ2 may be a positive regulator of VEGFA-induced microvascular permeability however this function appears to be specific to VEGFA, at least in lung endothelial cells, and is not likely to pertain to other permeability inducing factors.

**Figure 2.**
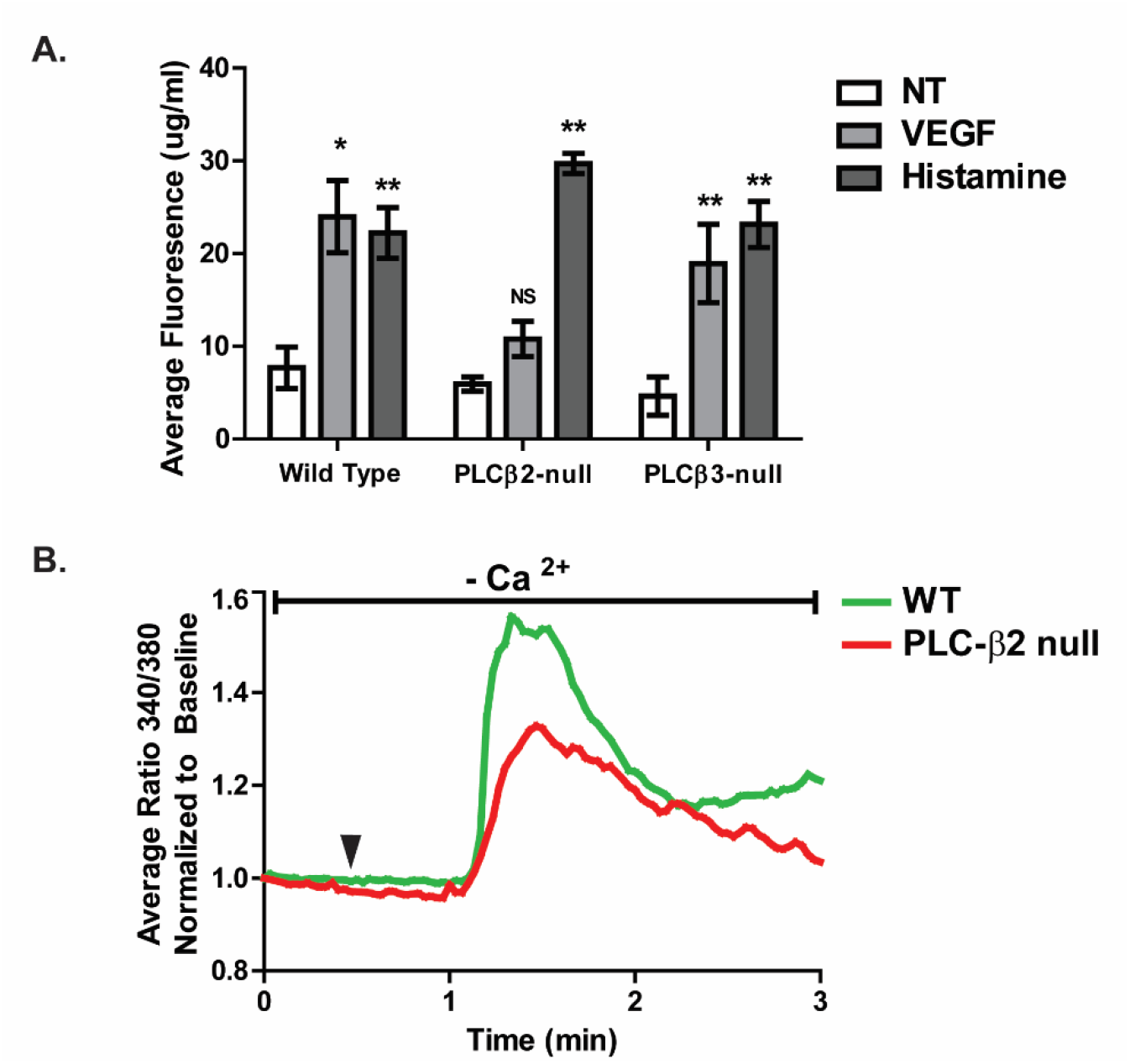
PLCβ2-null MLEC display a reduced VEGFA-induced permeability and calcium release in vitro. Primary lung endothelial cells were isolated from **wild type, PLCβ2-null** and **PLCβ3-null** animals and evaluated for endothelial permeability in a transwell assay. **A.** MLECs were treated with PBS control (**NT**), **VEGFA** or **histamine** and quantified for FITC in the lower chamber. (n = 3/treatment) Representative of at least three separate cell isolations and assays. **B. Wild type** (green) and **PLCβ2-null** (red) MLECs were treated with PBS control or VEGFA and intracellular calcium release was quantified (n = 10-15 individual cells/condition). Arrowhead indicates time of VEGFA treatment. (* = *P* ≤ 0.05; ** = *P* ≤ 0.01)

### PLCβ2 regulates endothelial cytosolic calcium flux in response to VEGFA

VEGFA binding to its receptor VEGFR2/KDR activates PLCγ and results in increased Ca^2+^ release from internal calcium stores as well as significant cytoplasmic calcium influx.^1,28,29^ Additionally, it has previously been shown that PLCβ3 is a negative regulator of VEGFA-induced calcium release in HUVECs and that loss of the PLCβ3 enzyme results in increased cytosolic calcium after treatment with VEGFA.^30^ In order to establish whether PLCβ2 can directly affect cytosolic calcium levels in endothelial cells downstream of VEGFA, murine lung endothelial cells (MLECs) were evaluated for intracellular calcium release following treatment with VEGFA. The isolated MLECs were imaged using Fura-2 labeling and treated with VEGFA in calcium free solution and multiple individual cells analyzed per sample (n=10-15 cells/condition). Intracellular calcium levels were rapidly increased in WT MLECs upon VEGFA treatment, Figure 2B. However, VEGFA treatment of PLCβ2-null MLECs resulted in a blunted intracellular calcium release that was reduced in amplitude by 50% when compared to WT MLECs. This result suggests that a suppressed calcium release after VEGFA binding to PLCβ2-null endothelial cells likely contributes to the reduced VEGFA-induced microvascular permeability observed in vivo.

In an attempt to establish a robust human cell model of the mouse primary cell system, human telomerase immortalized dermal microvascular endothelial (TIME) cells were used to evaluate VEGFA-induced total calcium flux. These cells were used to test whether calcium fluxes in human endothelial cells were similar to the observed responses in MLECs. VEGFA dose-response experiments recapitulated the expected increase in calcium levels in the TIME cells in a concentration and time dependent manner, Supplemental Figure 2A. To test the contribution of total PLC to the VEGFA-induced calcium flux, the global PLC inhibitor U73122 and an inactive analogue U73343 were used in a multi-well system to measure total calcium flux over time. As expected, U73122 completely blocks calcium increases after VEGFA treatment while the inactive analogue, U73343, had no significant effect when compared to DMSO control treatment, Supplemental Figure 2B.

In order to model the loss of either PLCβ2 or PLCβ3 in the transgenic murine models and specifically address the contribution of the PLCβ isoforms, the human TIME cells were stably transduced with lentiviral expressing shRNA constructs targeting PLCβ2 or PLCβ3 or GFP as a control. Effective suppression was observed by qRT-PCR (PLCβ2 – 77%, PLCβ3 – 87%) showing selective PLCβ isoform mRNA depletion with no evidence of compensation between PLCβ2 and PLCβ3, Figure 3A. The shTIME cells were then evaluated using real-time calcium imaging with Fura-2 dye loading followed by VEGFA stimulation in the presence of extracellular calcium. Figure 3B shows that PLCβ2 shTIME cells have a significantly reduced cytosolic calcium increase in response to VEGFA when compared to the shGFP control cells, which recapitulated the observations in the primary PLCβ2-null MLECs. PLCβ3 shTIME cells display a significantly increased flux in cytosolic calcium consistent with previous observations in mouse primary cells.^30^ Importantly, shGFP TIME cells displayed no change in VEGFA response profile when compared to parental TIME cells, Supplemental Figure 2A & Figure 3B. To define whether the regulation of cytosolic calcium flux by PLCβ isoforms observed was specific to VEGFA, histamine was employed as an alternative endothelial permeability activator. Contrary to VEGFA, histamine treatment resulted in a rapid increase in endothelial cell intracellular calcium release in the TIME cells. There was no difference in the calcium flux between the shGFP, shPLCβ2 and the shPLCβ3 cell lines, Figure 3C.

**Figure 3.**
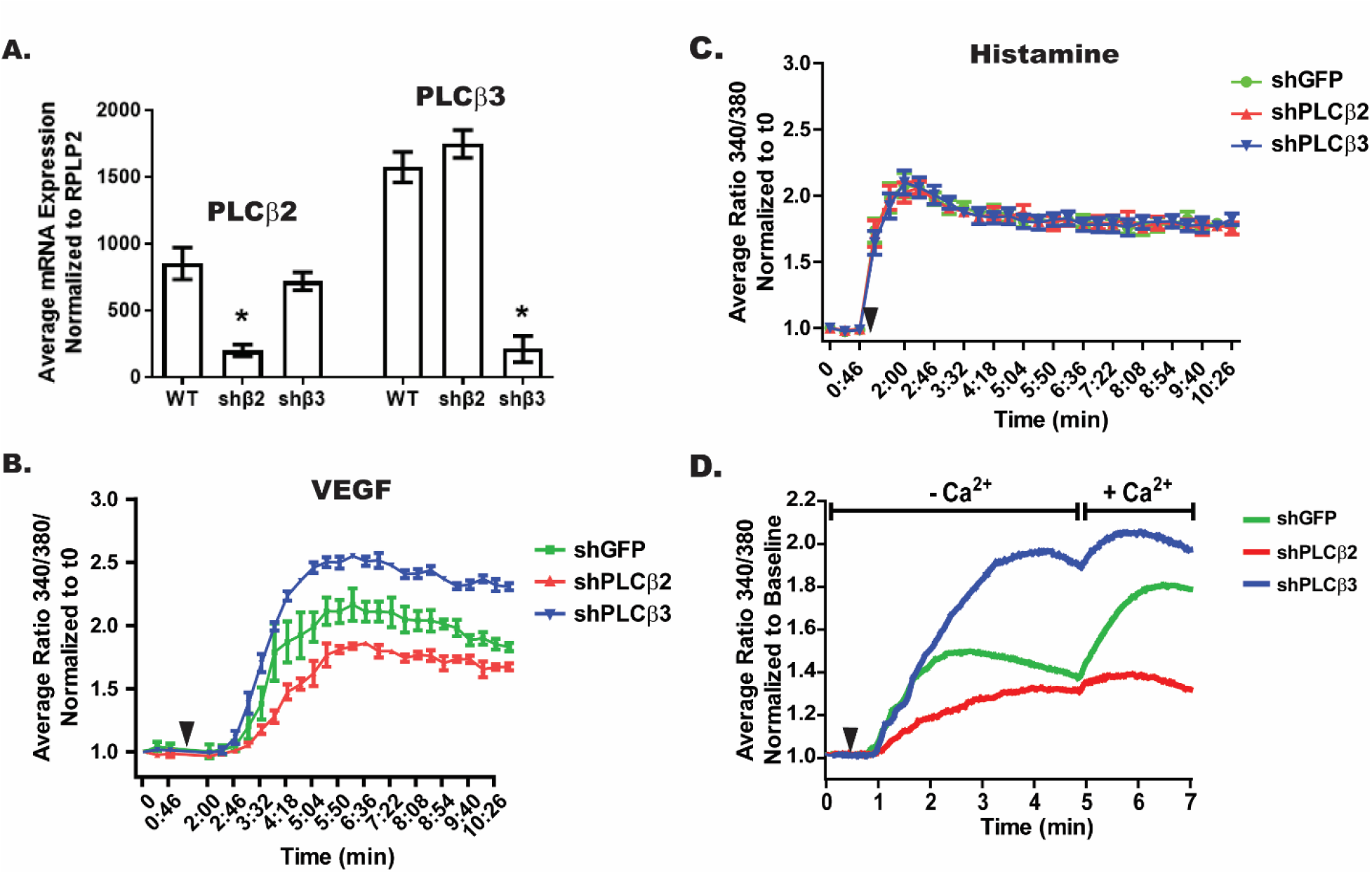
TIME shPLCβ2 cells display reduced total calcium flux after treatment with VEGFA. TIME control and PLCβ modified cells were established with **shGFP**, **shPLCβ2** and **shPLCβ3** lentiviral constructs and antibiotic selection. **A.** Target specific knock-down was verified via qRT-PCR for PLCβ2 and PLCβ3 mRNA levels. **B-C. shGFP** (green), **shPLCβ2** (red) and **shPLCβ3** (blue) TIME cells were assayed in a 96-well calcium flux assay following treatment with VEGFA or histamine. **D. shGFP** (green), **shPLCβ2** (red) and **shPLCβ3** (blue) TIME cells were evaluated in an intracellular calcium release (no extracellular calcium) and extracellular calcium entry assay (extracellular calcium). (n ≥ 24 individual cells/condition). Arrowhead indicates time of VEGFA/histamine treatment. (* = *P* ≤ 0.05)

To further dissect the change in calcium flux in these endothelial cells, individual cells were evaluated for VEGFA-induced intracellular calcium release (no extracellular calcium present) followed by VEGFA-induced extracellular calcium entry (extracellular calcium available) in a calcium release/entry imaging assay. Figure 3D shows the average of multiple cells from each cell line (n≥24 cells/sample) treated with VEGFA and evaluated for intracellular calcium release followed by extracellular calcium influx. PLCβ2 shTIME cells displayed significantly reduced initial intracellular calcium release as well as reduced calcium influx in response to VEGFA when compared to the shGFP control cells and PLCβ3 shRNA TIME cells display a significantly increased calcium levels both during intracellular calcium release and calcium entry. Taken together, endothelial cells with limited or no PLCβ2 present are defective at both intracellular calcium release and extracellular calcium entry into the cell subsequent to VEGFA stimulation and further support that PLCβ2 may be functioning as a positive regulator of VEGFA-induced cytosolic calcium flux.

### PLCβ2 regulates PIP2 levels in endothelial cells

PLC enzymes function in the cell membrane to cleave PIP2 into IP3 and DAG downstream of various stimuli, including VEGFA. These events result in changes in cytoplasmic calcium flux through IP3-induced calcium release of intracellular calcium from endoplasmic reticulum. As well as direct and indirect effects of DAG at the plasma membrane resulting in extracellular calcium entry into the cell as well as other signaling events.^25,26,42–44^ In order to assess events upstream of the observed differences in VEGFA-induced calcium flux, the cellular levels of PIP2 were evaluated in shTIME cells under basal and VEGFA-stimulated conditions.

Basal PIP2 levels were determined via ELISA in control or PLC beta modulated TIME cells, Figure 4A. shPLCβ2 cells were observed to have lower basal levels of PIP2 compared to shGFP and shPLCβ3 cells. No significant difference in PIP2 basal levels were observed in shPLCβ3 cells when compared to control cells. Importantly, upon stimulation with VEGFA, a rapid and significant decrease followed by restoration of the PIP2 level was observed in shGFP control cells, Figure 4B. However, there was no change in the low level of PIP2 detected in shPLCβ2 cells stimulated with VEGFA while shPLCβ3 cells displayed consistent elevated levels of PIP2 with no change during VEGFA stimulation. This data suggests that PLCβ2 regulates processes upstream of calcium flux and may directly contribute to the maintenance of cellular PIP2 levels and thus both PLCβ2 and PLCβ3 regulating the balance of PIP2.

**Figure 4.**
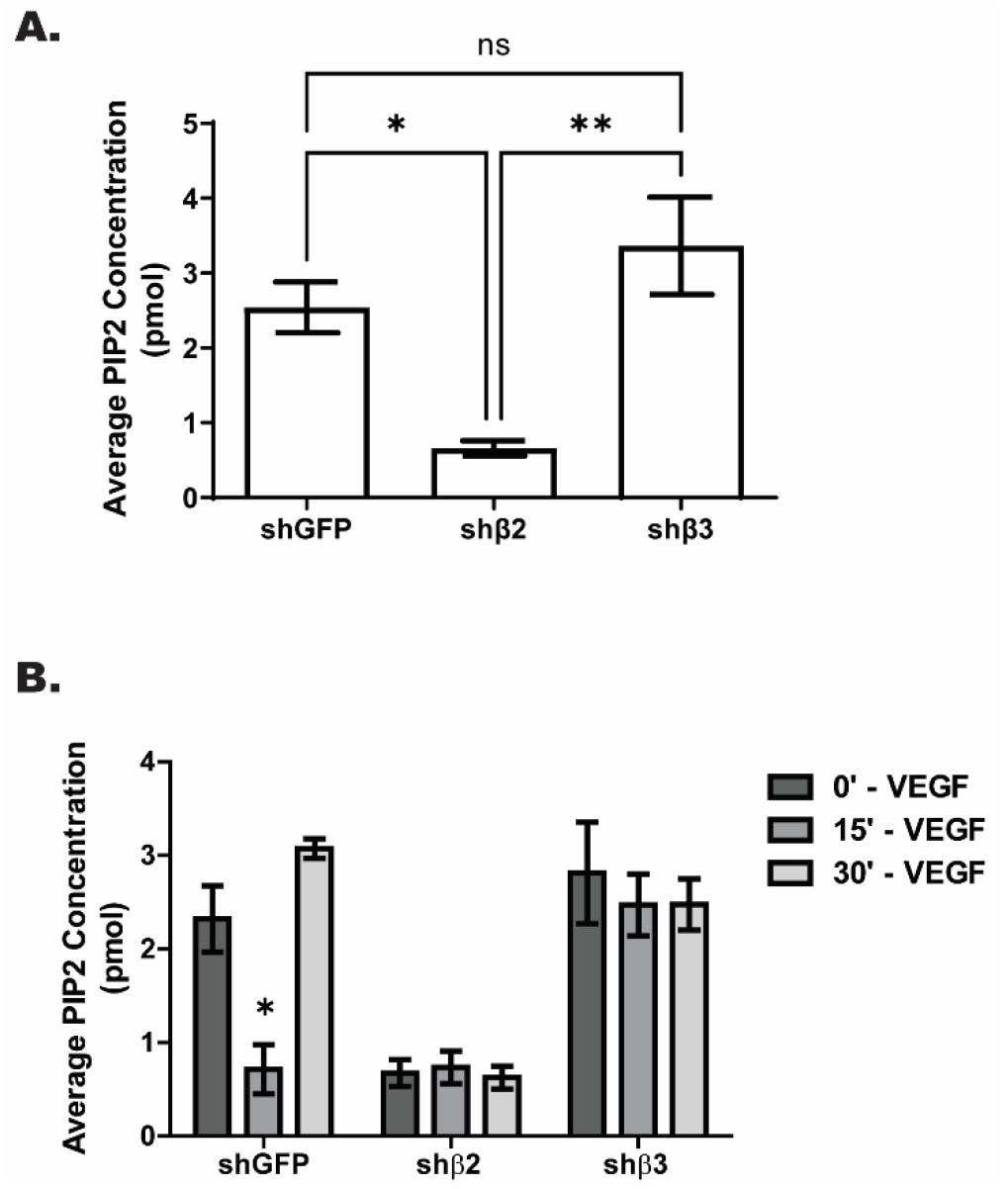
Basal and VEGFA stimulated PIP2 levels are reduced in TIME cells with modulated PLCβ2 expression. **A.** Basal PIP2 levels were quantified in **shGFP**, shPLCβ2 (**shβ2**) and shPLCβ3 (**shβ3**). (n = 3/condition) B. PIP2 levels following treatment with VEGFA were quantified at time 0-, 15- and 30-mins post treatment in **shGFP**, shPLCβ2 (**shβ2**) and **shPLCβ3** (**shβ3**) TIME cells. (n = 3/condition) (* = *P* ≤ 0.05; ** = *P* ≤ 0.01)

### PLCβ2-null mice are protected from injury in lung and heart models of vascular permeability

To determine whether these mechanisms of control by PLCβ2 in microvascular permeability would provide any benefit under pathological conditions, two disease models with a vascular permeability component were evaluated in WT and PLCβ2-null mice as a validation of the observed dermal vascular permeability response.

First, a model of lung oxygen injury was employed. This model aims to replicate conditions that are associated with acute lung injury (ALI) or acute respiratory distress syndrome (ARDS), both pathological conditions with minimal treatment options.^45,46^ It has been shown in various preclinical model species that exposure to hyperoxia results in changes in VEGFA levels in lung tissue. An initial increase in VEGFA in bronchoalveolar lavage (BAL) fluid followed by decreased VEGFA levels in persistent hyperoxic conditions.^47–49^ Upon return to room air, a dramatic increase in VEGFA levels is observed.^50,51^ In this study, animals were exposed to hyperoxia for 72 hours, followed by return to room air for 4 hours. WT mice exposed to these conditions displayed significant weight loss, increased lung permeability and changes in lung tissue morphology upon return to normal conditions. Importantly, when PLCβ2-null animals were challenged in this model, it was observed that the PLCβ2-null animals displayed a significantly reduced body weight loss when compared to WT animals after three days of hyperoxia, Figure 5. PLCβ2-null animals weight loss was approximately one-third of that observed for WT animals. Prior to harvest, animals were perfused via intravenous injection with FITC-dextran to evaluate the extent of vascular permeability in the lungs. BAL fluid was collected, and fluorescence measured. Wild-type animals demonstrated a 2.5-fold greater induction of lung permeability compared to the PLCβ2-null animals as measured by FITC-dextran in BAL fluid. Consistent with the observed permeability levels, histological analysis demonstrated not only air space or alveolar disruption (as measured by mean cord length) but peri-arteriolar edema was also markedly reduced in the PLCβ2-null group compared to the WT control group, 36% and 31% reduction respectively. Taken together these data, suggest that the PLCβ2-null animals may be protected from hyperoxia-induced lung injury and edema, through a direct reduction in pathological microvascular permeability.

**Figure 5.**
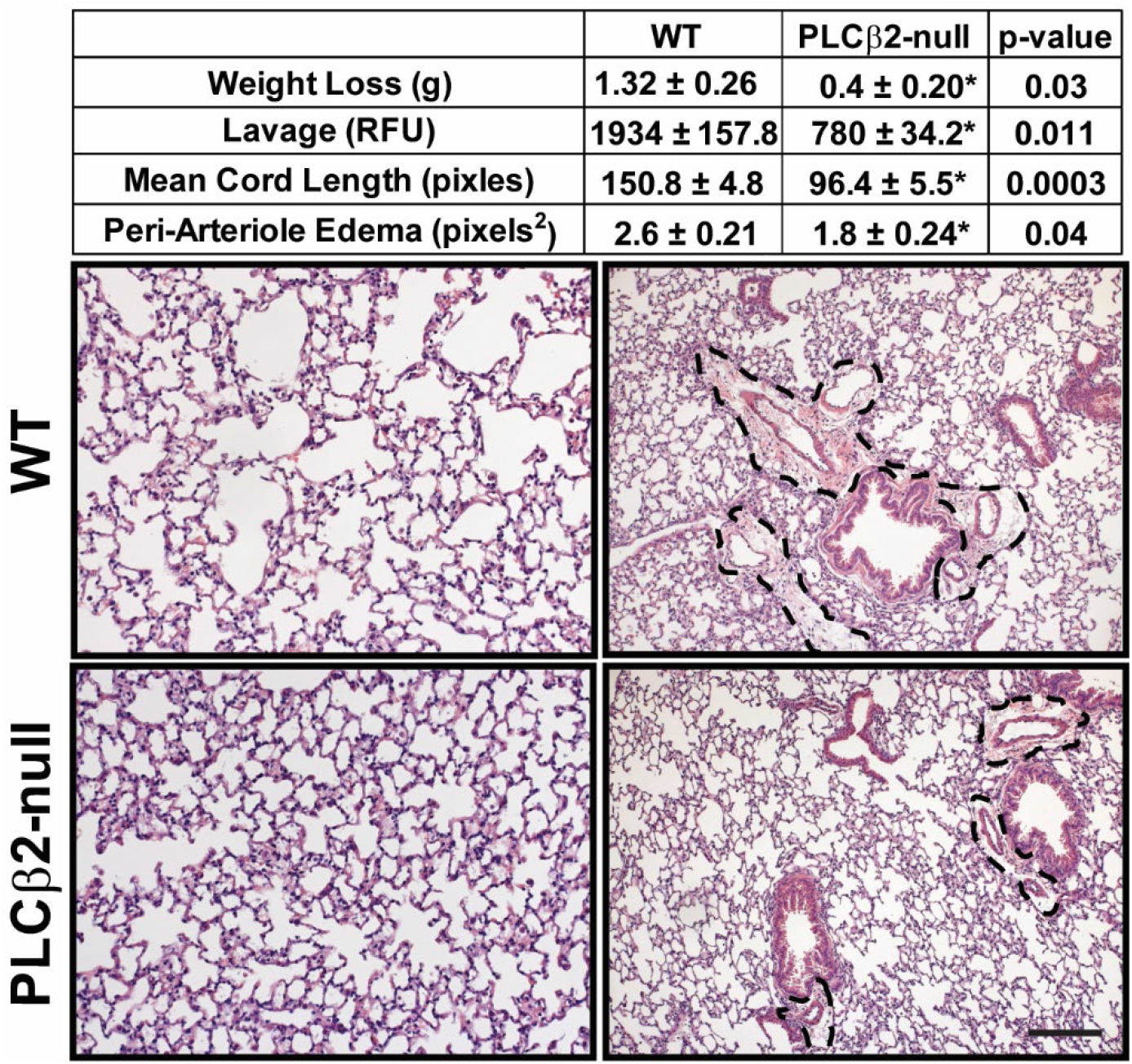
PLCβ2-null animals are resistant to hyperoxia-induced lung damage. **Wild-Type** and **PLCβ2-null** animals were exposed to hyperoxic conditions for 72hrs then moved to normoxic (room air) conditions for 4hrs. Body weight (g) (change from time 0), lung fluid lavage (vessel permeability) (RFU) and tissue histology end points (mean cord length and peri-arteriole edema) were quantified. Representative images of **WT** and **PLCβ2-null** tissues are shown. Bar = 100uM. Dotted lines show areas of edema. (n = 4/condition)

A second model was performed to evaluate the effect of PLCβ2 suppression in an in vivo disease model. An acute surgical model of cardiac ischemia:reperfusion (I:R) was performed on WT and PLCβ2-null mice. Previous work has shown that microvascular permeability and VEGFA are both increased following cardiac I:R injury across species and I:R injury in various other tissues; in fact, targeting VEGFA post-I:R has been proposed as a treatment to reduce tissue damage.^18,52–55^ In WT animals, a clear increase in vascular permeability is observed below the ligation site compared to sham treated animals, Figure 6A. Similar to the acute lung injury model, the hearts of PLCβ2-null mice appear to be protected and display less permeability in the region directly below the site of LAD when compared to WT controls, Figure 6B. Additional animals in the same model were allowed to recover for 10 weeks prior to euthanasia and tissue was analyzed in order to assess heart damage. When the heart tissues were processed and analyzed for infarct size via trichrome staining, Figure 6C, it was found that the PLCβ2-null animals had 37.8% smaller infarct region compared to the WT control animals, Figure 6D. This data suggests that the decreased vascular permeability immediately following cardiac I:R injury in PLCβ2-null mice, may lead to reduced permanent heart damage similar to the protection observed in the hyperoxia lung model.

**Figure 6.**
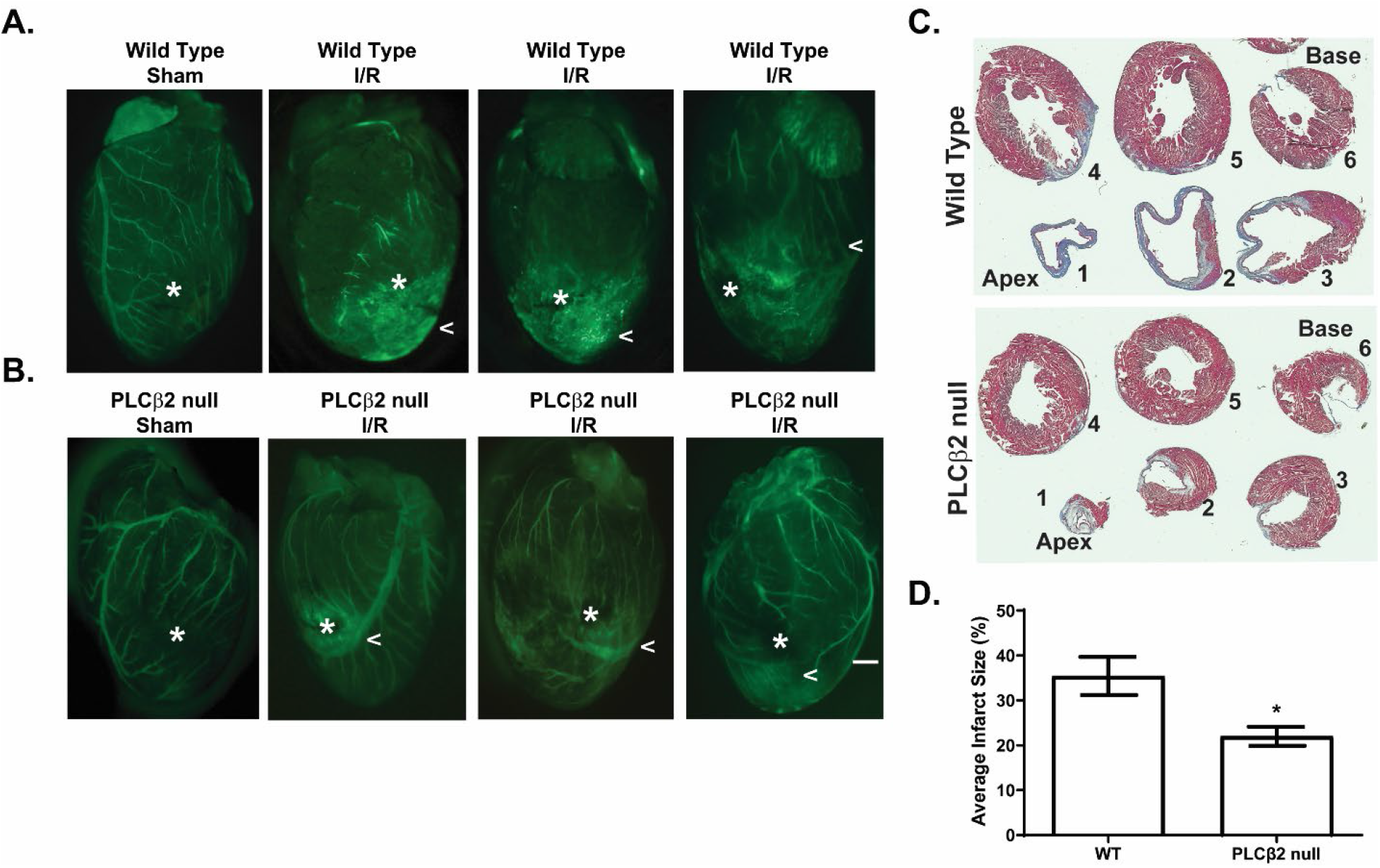
PLCβ2-null animals are protected from cardiac ischemia:reperfusion damage. **Wild-Type** and **PLCβ2-null** animals were evaluated in a cardiac ischemia:reperfusion model of LAD ligation. **A.** Representative images gross tissue permeability of wild type (**WT**) control (**Sham**) or treated (**I:R**). Asterix indicates the site of LAD. Carrot highlights areas of increased permeability. **B.** Representative images of **PLCβ2-null** control (**Sham**) or treated (**I:R**) following 30mins of I:R. Asterix indicates the site of LAD. Carrot highlights areas of increased permeability. Bar = 1 mm. **C.** Representative images from **Wild-Type** and **PLCβ2-null** trichrome stained tissue sections from the apex (1) to the base (6) of the heart. **D.** Quantification of heathy tissue (red) and infarct region (blue) normalized to percent total tissue. (n = 4-6/group) (* = *P* ≤ 0.05)

## Discussion

Here, we identify PLCβ2 as a novel, potent inducer of VEGFA-mediated VP, and we demonstrate that suppression of PLCβ2 can limit maximal VEGFA-induced endothelial and vessel permeability. As previously reported, PLCβ3-null animals display hypersensitivity to VEGFA-induced microvascular permeability when compared to WT counterparts, supporting the data that suggest the PLCβ3 is a negative regulator of VEGFA-induced VP.^30^ However, when PLCβ2-null animals were evaluated, these mice demonstrated no detectable increase in VP after VEGFA exposure, suggesting that PLCβ2 is actually a positive regulator of VEGFA-induced VP and that these mice could possibly be resistant to pathologically induced VP triggered by VEGFA. Importantly, there was no evidence of differences in baseline homeostatic permeability in PLCβ2-null or PLCβ3-null when compared to WT controls. While increased permeability by dermal VP assay was initially observed in the vehicle control group of PLCβ3-null mice compared to WT controls, no significant differences were observed between WT and PLCβ2-null or PLCβ3-null animals in peripheral basal vascular permeability in any of the eight organs tested.

Importantly, and consistent with other PLC signaling following VEGFA treatment, the suppression of VP in PLCβ2-null or shRNA models, occurs at the initial amplitude of calcium release from intracellular stores, and, from pronounced reduction of entry from extracellular stores. These data indicate that VEGFA activation cascades are likely to promote cell membrane calcium channels to function for an extended time promoting the intracellular calcium entry and maintaining the calcium levels necessary for the observed junctional or transcellular events in VEGFA-induced microvascular permeability. In addition, the PLC-beta regulation of cytosolic calcium appears to be preferential to VEGFA stimulation since suppression or elimination of PLCβ2 had no effect on histamine-induced acute calcium release downstream of the H1 GPCR and PLCβ activation. These data clearly imply that the VEGFR2 signaling by VEGFA is likely restricted to its own membrane clustering domain and does not affect other PLC-signaling receptors such as H1 GPCR.

This is the first report of regulation of VEGF signaling in endothelial cells by PLCβ2. However, this is not the first report of this type of regulation by PLCβ2. A similar effect has been shown in neutrophils lacking the PLCβ2 isoform where both IP3 production and calcium flux were reduced in response to chemoattractant stimulus.^32^ Additionally it has been demonstrated that in sites of ischemia, endothelial progenitor cell recruitment is regulated through PLCβ2 downstream of insulin-like growth factor 2 stimulation by regulation of intracellular calcium flux.^56^

PIP2 is a critical regulator downstream of PLC activation. The data presented here suggest that the dynamics of PIP2 levels in endothelial cells changes dramatically based on whether the PLCβ isoform is predominantly β3 or β2. Clearly in the TIME cells lacking PLCβ2, the levels of total cellular PIP2 are suppressed and do not change with VEGFA stimulation. Conversely, when PLCβ3 is lacking, the PIP2 levels appear to be maintained even during VEGFA-activation and significant increased calcium flux leading to VP. The mechanism of these competitive dynamics is unclear. One possibility is the clustering of selective PLCβ isoforms within membrane compartments, or that the PLCβ3 isoform is preferentially maintained at the cell membrane/GPCR association thus providing baseline PIP2 cleavage. This complex mechanism may also include the PIP and PIP2 kinases and their regulation via upstream or downstream components and will require further investigation. It is possible that changes in PIP2 were not observed in PLCβ2 or PLCβ3 knockdown TIME cells due to a limit of resolution in the assay. Additional time points may reveal an alteration in the timing of the PIP2 dynamics in these cells following treatment with VEGFA. However, it is clear that PLCβ3 negatively regulates VEGFA-induced vascular permeability ^30^, whereas PLCβ2 promotes endothelial permeability mediated by VEGFA downstream of both PIP2 and calcium effects.

Gaining a greater understanding of how concomitant regulation of different PLC isozymes controls cellular function may have important clinical implications. For example, PLC is one of the few signaling effectors that is directly regulated by opioids. Chronic morphine treatment regulates PLC in an isozyme-specific manner by causing increased phosphorylation of PLCβ3 and reduced phosphorylation of PLCβ1. This shifts the relative balance of isoform activity in favor of PLCβ1 because both of these PLCs are negatively modulated by phosphorylation.^57^ The effects of morphine and other opioids on the activity of PLCβ2 remain unknown. Like morphine, anticancer drugs designed to inhibit G-protein signaling also reduce PLCβ3 activity.^58,59^ Given that PLCβ3 has been shown to negatively regulate VEGFA-mediated vascular permeability,^30^ drugs that inhibit PLCβ3 activity have the potential to increase vascular permeability and elevate the risk of subsequent edema and tissue damage. Our work presented here suggests inhibition of PLCβ2 could be a viable therapeutic strategy to counteract such increases in vascular permeability.

The possibility exists that a selective inhibitor to PLCβ2 maybe a highly effective strategy for treating cardiac and stroke reperfusion as well as lung damage associated with chemical or ventilator-induced suppression of oxygen transfer. While there have been several reports of various pan PLC inhibitors^60–63^, the ability to develop isoform specific inhibitors, particularly for the closely related beta isoforms is, and will likely continue to be, a difficult proposition. This challenge stems from the fact that PLC isozymes share a conserved binding pocket and mechanism of catalysis, and the small molecule inhibitors that have been developed bind to the active site that is shared among isozymes.^60^ Given these challenges with small molecule inhibitors, investigation of alternative molecular inhibitors will be required. In summary, the dynamic and relative balance of PLC isozymes regulates endothelial cell function, and in particular, the PLCβ2 isoform promotes vascular permeability, whereas PLCβ3 protects endothelial barrier integrity.

## Authorship

Contribution: KNP, ZY, and CGC performed experiments; KNP, ZY, LY, CGC, BTL, LHH and KPC contributed to discussions on the design of the experiments and/or interpretation of the data; KNP wrote the manuscript with contributions and comments from LHH and KPC.

## Conflict-of-interest disclosure

The authors declare no competing financial interests.

## Acknowledgements

The authors would like to thank Dianqing Wu, PhD (Yale School of Medicine, New Haven, CT) for the PLCβ2-null and PLCβ3-null mice used in this study and the University of Connecticut Health Center Research Histology Core for assistance with tissue processing and sectioning.

This work was supported by funding to KPC from the NIH (2P01 HL070694-06A1) and the CT Department of Public Health. The contributions of LHH were supported by funding from American Cancer Society (RSG-21-034-01-TBG), Windfeldt Cancer Research Award, and the Hormel Foundation.

## Nonstandard Abbreviations and Acronyms

BAL: bronchoalveolar lavage
DAG: diacylglycerol
FGF-2: fibroblast growth factor 2
IP3: inositol 1,4,5-trisphosphate
I:R: ischemia:reperfusion
HIF: hypoxia-inducible factor
MLEC: murine lung endothelial cell
PIP2: phosphatidylinositol 4,5-bisphosphate
PLC: phospholipase C
TIME cells: telomerase-immortalized microvascular endothelial cells
TNF-alpha: tumor necrosis factor alpha
VEGF: vascular endothelial growth factor
VEGFR: vascular endothelial growth factor receptor
VP: vascular permeability
VPF: vascular permeability factor
VVO: vesiculo-vacuolar organelles
WT: wild type

## Notes

### Competing Interest Statement

The authors have declared no competing interest.

## References

1. Goddard LM, Iruela-Arispe ML. Cellular and molecular regulation of vascular permeability. Thrombosis and haemostasis. 2013;109(3):407–415.

2. Stevens T, Garcia JGN, Shasby DM, Bhattacharya J, Malik AB. Mechanisms regulating endothelial cell barrier function. American journal of physiology. Lung cellular and molecular physiology. 2000;279(3):419–L422.

3. Nagy J, Benjamin L, Zeng H, Dvorak A, Dvorak H. Vascular permeability, vascular hyperpermeability and angiogenesis. Angiogenesis (London). 2008;11(2):109–119.

4. Cheresh DA, Paul R, Zhang ZG, Eliceiri BP, Jiang Q, Boccia AD, Zhang RL, Chopp M. Src deficiency or blockade of Src activity in mice provides cerebral protection following stroke. Nature medicine. 2001;7(2):222–227.

5. Lionetti V, Recchia FA, Ranieri VM. Overview of ventilator-induced lung injury mechanisms. Current opinion in critical care. 2005;11(1):82–86.

6. Orfanos S, Mavrommati I, Korovesi I, Roussos C. Pulmonary endothelium in acute lung injury: from basic science to the critically ill. Intensive care medicine. 2004;30(9):1702–1714.

7. Senger DR, Galli SJ, Dvorak AM, Perruzzi CA, Harvey VS, Dvorak HF. Tumor cells secrete a vascular permeability factor that promotes accumulation of ascites fluid. Science. 1983;219(4587):983–985.

8. Senger DR, Water LVD, Brown LF, Nagy JA, Yeo K-T, Yeo T-K, Berse B, Jackman RW, Dvorak AM, Dvorak HF. Vascular permeability factor (VPF, VEGF) in tumor biology. Cancer and Metastasis Reviews. 1993;12(3–4):303–324.

9. Detmar M, Brown LF, Berse B, Jackman RW, Elicker BM, Dvorak HF, Claffey KP. Hypoxia Regulates the Expression of Vascular Permeability Factor/Vascular Endothelial Growth Factor (VPF/VEGF) and its Receptors in Human Skin. Journal of Investigative Dermatology. 1997;108(3):263–268.

10. Dvorak HF. Reconciling VEGF With VPF: The Importance of Increased Vascular Permeability for Stroma Formation in Tumors, Healing Wounds, and Chronic Inflammation. Frontiers in Cell and Developmental Biology. 2021;9:660609.

11. Bates DO, Harper SJ. Regulation of vascular permeability by vascular endothelial growth factors. Vascular Pharmacology. 2002;39(4):225–237.

12. Ferrara N, Gerber H-P, LeCouter J. The biology of VEGF and its receptors. Nature medicine. 2003;9(6):669–676.

13. Bates DO. Vascular endothelial growth factors and vascular permeability. Cardiovascular Research. 2010;87(2):262–271.

14. Ziyad S, Iruela-Arispe ML. Molecular Mechanisms of Tumor Angiogenesis. Genes & Cancer. 2011;2(12):1085–1096.

15. Apte RS, Chen DS, Ferrara N. VEGF in Signaling and Disease: Beyond Discovery and Development. Cell. 2019;176(6):1248–1264.

16. Seghezzi G, Patel S, Ren CJ, Gualandris A, Pintucci G, Robbins ES, Shapiro RL, Galloway AC, Rifkin DB, Mignatti P. Fibroblast Growth Factor-2 (FGF-2) Induces Vascular Endothelial Growth Factor (VEGF) Expression in the Endothelial Cells of Forming Capillaries: An Autocrine Mechanism Contributing to Angiogenesis. The Journal of cell biology. 1998;141(7):1659–1673.

17. Ushiro S, Ono M, Izumi H, Kohno K, Taniguchi N, Higashiyama S, Kuwano M. Heparin-binding epidermal growth factor-like growth factor: p91 activation induction of plasminogen activator/inhibitor, and tubular morphogenesis in human microvascular endothelial cells. Japanese journal of cancer research (Gann). 1996;87(1):68–77.

18. Crafts TD, Jensen AR, Blocher-Smith EC, Markel TA. Vascular endothelial growth factor: Therapeutic possibilities and challenges for the treatment of ischemia. Cytokine (Philadelphia, Pa.). 2015;71(2):385–393.

19. Forsythe JA, Jiang BH, Iyer NV, Agani F, Leung SW, Koos RD, Semenza GL. Activation of vascular endothelial growth factor gene transcription by hypoxia-inducible factor 1. Molecular and Cellular Biology. 1996;16(9):4604–4613.

20. Dvorak AM, Feng D. The Vesiculo–Vacuolar Organelle (VVO): A New Endothelial Cell Permeability Organelle. Journal of Histochemistry & Cytochemistry. 2000;49(4):419–431.

21. Simons M, Gordon E, Claesson-Welsh L. Mechanisms and regulation of endothelial VEGF receptor signalling. Nature reviews. Molecular cell biology. 2016;17(10):611–625.

22. Claesson-Welsh L, Olsson A-K, Dimberg A, Kreuger J. VEGF receptor signalling ? in control of vascular function. Nature reviews. Molecular cell biology. 2006;7(5):359–371.

23. Abhinand CS, Raju R, Soumya SJ, Arya PS, Sudhakaran PR. VEGF-A/VEGFR2 signaling network in endothelial cells relevant to angiogenesis. Journal of Cell Communication and Signaling. 2016;10(4):347–354.

24. Katan M, Cockcroft S. Phospholipase C families: Common themes and versatility in physiology and pathology. Progress in Lipid Research. 2020;80:101065.

25. Bunney TD, Katan M. PLC regulation: emerging pictures for molecular mechanisms. Trends in Biochemical Sciences. 2011;36(2):88–96.

26. Yang YR, Jang H-J, Ryu SH, Suh P-G. Phospholipases in Health and Disease. 2014:3–38.

27. Chen D, Simons M. Emerging roles of PLCγ1 in endothelial biology. Science Signaling. 2021;14(694).

28. Mukhopadhyay D, Zeng H. Involvement of G Proteins in Vascular Permeability Factor/ Vascular Endothelial Growth Factor Signaling. Cold Spring Harbor Symposia on Quantitative Biology. 2002;67(0):275–284.

29. Wu H mac, Yuan Y, Zawieja DC, Tinsley J, Granger HJ. Role of phospholipase C, protein kinase C, and calcium in VEGF-induced venular hyperpermeability. American Journal of Physiology-Heart and Circulatory Physiology. 1999;276(2):H535–H542.

30. Hoeppner LH, Phoenix KN, Clark KJ, Bhattacharya R, Gong X, Sciuto TE, Vohra P, Suresh S, Bhattacharya S, Dvorak AM, Ekker SC, Dvorak HF, Claffey KP, Mukhopadhyay D. Revealing the role of phospholipase Cβ3 in the regulation of VEGF-induced vascular permeability. Blood. 2012;120(11):2167–2173.

31. Xie W, Samoriski GM, McLaughlin JP, Romoser VA, Smrcka A, Hinkle PM, Bidlack JM, Gross RA, Jiang H, Wu D. Genetic alteration of phospholipase C β3 expression modulates behavioral and cellular responses to μ opioids. Proceedings of the National Academy of Sciences. 1999;96(18):10385–10390.

32. Li Z, Jiang H, Xie W, Zhang Z, Smrcka AV, Wu D. Roles of PLC-beta2 and -beta3 and PI3Kgamma in chemoattractant-mediated signal transduction. Science (American Association for the Advancement of Science). 2000;287(5455):1046–1049.

33. Sanchez T, Estrada-Hernandez T, Paik J-H, Wu M-T, Venkataraman K, Brinkmann V, Claffey K, Hla T. Phosphorylation and Action of the Immunomodulator FTY720 Inhibits Vascular Endothelial Cell Growth Factor-induced Vascular Permeability. The Journal of biological chemistry. 2003;278(47):47281–47290.

34. Wick MJ, Harral JW, Loomis ZL, Dempsey EC. An Optimized Evans Blue Protocol to Assess Vascular Leak in the Mouse. Journal of Visualized Experiments. 2018;(139).

35. Su L-T, Agapito MA, Li M, Simonson WTN, Huttenlocher A, Habas R, Yue L, Runnels LW. TRPM7 Regulates Cell Adhesion by Controlling the Calcium-dependent Protease Calpain. The Journal of biological chemistry. 2006;281(16):11260–11270.

36. Sonin D, Zhou S-Y, Cronin C, Sonina T, Wu J, Jacobson KA, Pappano A, Liang BT. Role of P2X purinergic receptors in the rescue of ischemic heart failure. American Journal of Physiology-Heart and Circulatory Physiology. 2008;295(3):H1191–H1197.

37. Pereira FE, Cronin C, Ghosh M, Zhou S-Y, Agosto M, Subramani J, Wang R, Shen J-B, Schacke W, Liang B, Yang TH, McAulliffe B, Liang BT, Shapiro LH. CD13 is essential for inflammatory trafficking and infarct healing following permanent coronary artery occlusion in mice. Cardiovascular Research. 2013;100(1):74–83.

38. Chen L, Chen CX, Gan XT, Beier N, Scholz W, Karmazyn M. Inhibition and reversal of myocardial infarction-induced hypertrophy and heart failure by NHE-1 inhibition. American Journal of Physiology-Heart and Circulatory Physiology. 2004;286(1):H381–H387.

39. Lorenzo AD, Fernández-Hernando C, Cirino G, Sessa WC. Akt1 is critical for acute inflammation and histamine-mediated vascular leakage. Proceedings of the National Academy of Sciences. 2009;106(34):14552–14557.

40. Moy AB, Engelenhoven JV, Bodmer J, Kamath J, Keese C, Giaever I, Shasby S, Shasby DM. Histamine and thrombin modulate endothelial focal adhesion through centripetal and centrifugal forces. Journal of Clinical Investigation. 1996;97(4):1020–1027.

41. Ashina K, Tsubosaka Y, Nakamura T, Omori K, Kobayashi K, Hori M, Ozaki H, Murata T. Histamine Induces Vascular Hyperpermeability by Increasing Blood Flow and Endothelial Barrier Disruption In Vivo. PLoS ONE. 2015;10(7):e0132367.

42. Sekiya F. Encyclopedia of Biological Chemistry (Second Edition). Signaling: Article Titles: P. 2013;(Experimental Cell Research2531999):467–471.

43. Béziau DM, Toussaint F, Blanchette A, Dayeh NR, Charbel C, Tardif J-C, Dupuis J, Ledoux J. Expression of Phosphoinositide-Specific Phospholipase C Isoforms in Native Endothelial Cells. PloS one. 2015;10(4):e0123769.

44. Cocco L, Follo MY, Manzoli L, Suh P-G. Phosphoinositide-specific phospholipase C in health and disease. Journal of Lipid Research. 2015;56(10):1853–1860.

45. Radu M, Chernoff J. An in vivo Assay to Test Blood Vessel Permeability. Journal of Visualized Experiments. 2013;(73):e50062.

46. Matute-Bello G, Frevert CW, Martin TR. Animal models of acute lung injury. American Journal of Physiology - Lung Cellular and Molecular Physiology. 2008;295(3):L379–L399.

47. Bhandari V. Hyperoxia-derived lung damage in preterm infants. Seminars in Fetal and Neonatal Medicine. 2010;15(4): 223–229.

48. Giusto K, Wanczyk H, Jensen T, Finck C. Hyperoxia-induced bronchopulmonary dysplasia: better models for better therapies. Disease Models & Mechanisms. 2021;14(2):dmm047753.

49. Mura M, Santos CC dos, Stewart D, Liu M. Vascular endothelial growth factor and related molecules in acute lung injury. Journal of Applied Physiology. 2004;97(5):1605–1617.

50. Watkins RH, D’Angio CT, Ryan RM, Patel A, Maniscalco WM. Differential expression of VEGF mRNA splice variants in newborn and adult hyperoxic lung injury. American Journal of Physiology-Lung Cellular and Molecular Physiology. 1999;276(5):L858–L867.

51. Maniscalco WM, Watkins RH, Finkelstein JN, Campbell MH. Vascular endothelial growth factor mRNA increases in alveolar epithelial cells during recovery from oxygen injury. American Journal of Respiratory Cell and Molecular Biology. 1995;13(4):377–386.

52. Neri M, Riezzo I, Pascale N, Pomara C, Turillazzi E. Ischemia/Reperfusion Injury following Acute Myocardial Infarction: A Critical Issue for Clinicians and Forensic Pathologists. Mediators of Inflammation. 2017;2017:7018393.

53. Anon. Myocardial capillary permeability after regional ischemia and reperfusion in the in vivo canine heart. Effect of superoxide dismutase.

54. Dobschuetz E von, Meyer S, Thorn D, Marme D, Hopt UT, Thomusch O. Targeting Vascular Endothelial Growth Factor Pathway Offers New Possibilities to Counteract Microvascular Disturbances During Ischemia/Reperfusion of the Pancreas. Transplantation. 2006;82(4):543–549.

55. Yla-Herttuala S, Rissanen TT, Vajanto I, Hartikainen J. Vascular Endothelial Growth Factors: Biology and Current Status of Clinical Applications in Cardiovascular Medicine. Journal of the American College of Cardiology. 2007;49(10):1015–1026.

56. Maeng Y-S, Choi H-J, Kwon J-Y, Park Y-W, Choi K-S, Min J-K, Kim Y-H, Suh P-G, Kang K-S, Won M-H, Kim Y-M, Kwon Y-G. Endothelial progenitor cell homing: prominent role of the IGF2-IGF2R-PLCβ2 axis. Blood. 2009;113(1):233–243.

57. Chakrabarti S, Liu N-J, Gintzler AR. Reciprocal modulation of phospholipase Cβ isoforms: Adaptation to chronic morphine. Proceedings of the National Academy of Sciences. 2003;100(23):13686–13691.

58. Prévost GP, Lonchampt MO, Holbeck S, Attoub S, Zaharevitz D, Alley M, Wright J, Brezak MC, Coulomb H, Savola A, Huchet M, Chaumeron S, Nguyen Q-D, Forgez P, Bruyneel E, et al. Anticancer Activity of BIM-46174, a New Inhibitor of the Heterotrimeric Gα/Gβγ Protein Complex. Cancer Research. 2006;66(18):9227–9234.

59. Guha S, Eibl G, Kisfalvi K, Fan RS, Burdick M, Reber H, Hines OJ, Strieter R, Rozengurt E. Broad-Spectrum G Protein–Coupled Receptor Antagonist, [D-Arg1,D-Trp5,7,9,Leu11]SP: A Dual Inhibitor of Growth and Angiogenesis in Pancreatic Cancer. Cancer Research. 2005;65(7):2738–2745.

60. Huang W, Barrett M, Hajicek N, Hicks S, Harden TK, Sondek J, Zhang Q. Small Molecule Inhibitors of Phospholipase C from a Novel High-throughput Screen*. Journal of Biological Chemistry. 2013;288(8):5840–5848.

61. Wong R, Fabian L, Forer A, Brill JA. Phospholipase C and myosin light chain kinase inhibition define a common step in actin regulation during cytokinesis. BMC Cell Biology. 2007;8(1):15.

62. Rees SWP, Leung E, Reynisson J, Barker D, Pilkington LI. Development of 2-Morpholino-N-hydroxybenzamides as anti-proliferative PC-PLC inhibitors. Bioorganic Chemistry. 2021;114:105152.

63. Oh WK, Oh H, Kim BY, Kim BS, Ahn JS. CRM-51006, a New Phospholipase C (PLC) Inhibitor, Produced by Unidentified Fungal Strain MT51005. The Journal of Antibiotics. 2004;57(12):808–811.

