## Supplemental Figures for "Phospholipase Cβ2 Promotes Vascular Endothelial Growth Factor Induced Vascular Permeability"

**Supplemental Figure 1**

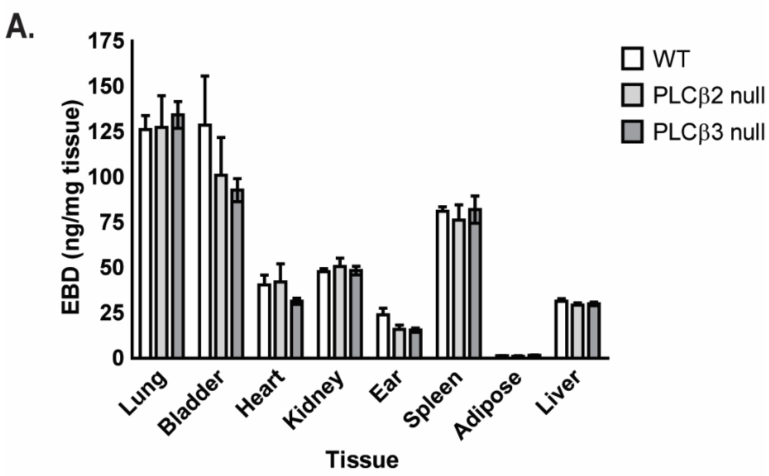


**Supplemental Figure 1 - Basal Permeability in WT, PLCβ2-null and PLCβ3-null tissues. Wild-Type** (white bars), **PLCβ2-null** (light grey bars) and **PLCβ3-null** (dark grey bars) animals were injected IV with Evans's blue dye and tissues harvested after four hours. Quantification of dye in tissues is shown as ug of dye normalized to dry tissue weight (ng/mg). (n = 4/group)

**Supplemental Figure 2**

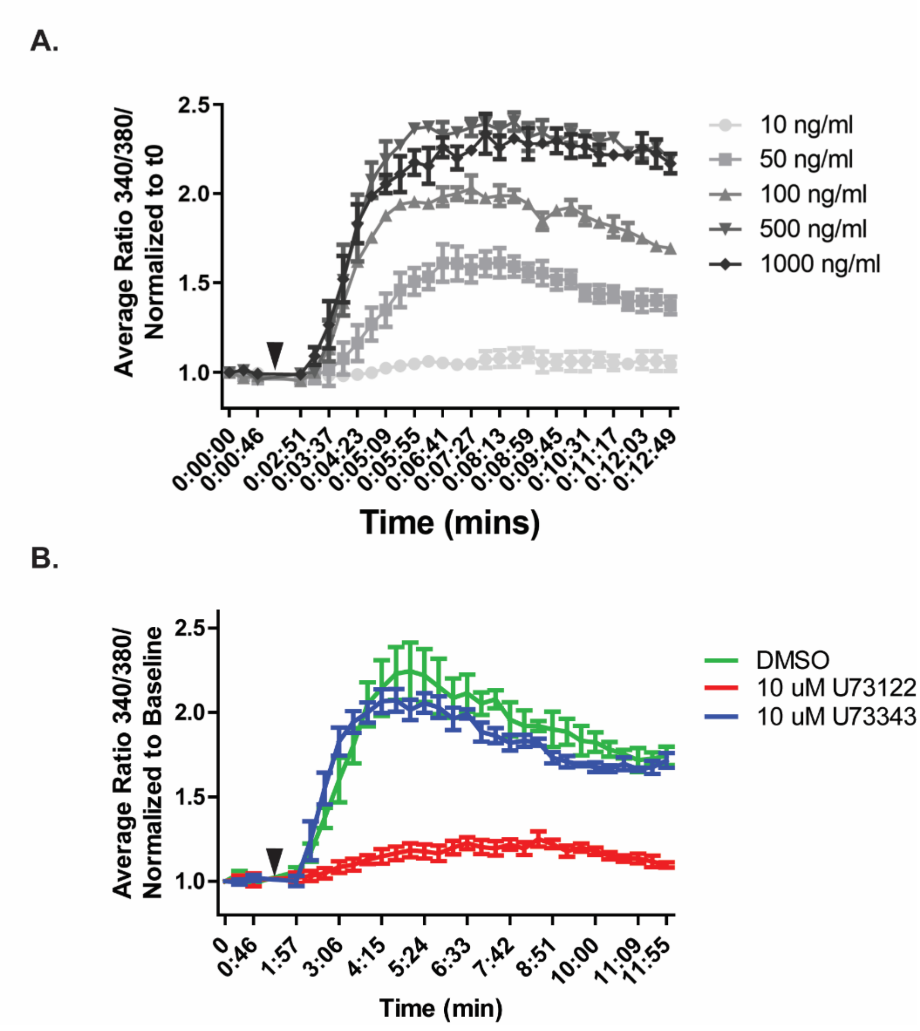


**Supplemental Figure 2 - PLC inhibition completely blocks VEGFA-induced calcium flux in TIME cells. A.** TIME cells were treated with increasing doses of VEGFA (10–1000 ng/ml) and calcium flux quantified over time. **B.** TIME cells were pre-treated with a pan-PLC inhibitor, **U73122** (red), an inactive analog, **U73313** (blue) or vehicle control, **DMSO** (green) then treated with VEGFA (100ng/ml). Calcium flux was quantified over time. Arrowhead indicates time of VEGFA treatment. (n = 4/dose)
